# Non-invasive real-time access to spatial attention information from 3T fMRI BOLD signals

**DOI:** 10.1101/2021.11.24.469873

**Authors:** S. Dali, C. Loriette, C. De Sousa, S. Clavagnier, F. Lamberton, D. Ibarrola, S. Ben Hamed

**Affiliations:** Institut des Sciences Cognitives Marc Jeannerod, CNRS UMR 5229, Université Claude Bernard Lyon 1, 67 Boulevard Pinel, 69675 Bron Cedex, France; CERMEP, CNRS UAR 2053, Lyon1 Université, Inserm US65, 59 Boulevard Pinel, 69677 Bron Cedex, France

**Keywords:** fMRI, decoding, spatial attention, machine learning, single subjects

## Abstract

Decoding internally generated cognitive states at the single-trial level remains challenging, particularly when these states are spatially specific and dynamically expressed over time. Here, we investigated whether the locus of covert spatial attention could be decoded from non-invasive 3T fMRI BOLD signals using a participant-specific pipeline designed for real-time implementation. Fourteen participants performed a spatially cued attention task in which attention was covertly directed toward one of eight possible locations while central fixation was maintained. Multivoxel patterns were classified using a linear support vector machine with leave-one-block-out cross-validation and feature selection performed independently within each training fold. Decoding accuracy substantially exceeded the theoretical chance level of 12.5%, reaching approximately 35–40% around target presentation and up to ∼60% in the best-performing participants. Classification errors were spatially structured, occurring preferentially at locations neighboring the cued position. Regional analyses showed that fine-grained spatial information was predominantly carried by occipital cortex, whereas parietal voxels showed a broader contralateral organization but limited eight-location decoding. Cross-temporal analyses revealed that the spatial representation generalized across successive BOLD samples, while trial-wise decoding suggested increasing concentration of the decoded attentional locus around the cued location as target onset approached. Decoding performance remained reproducible across four scanning sessions and improved with increasing amounts of training data. Together, these findings demonstrate that participant-specific 3T fMRI activity contains spatially precise and reproducible information about covert attention at the single-trial level, providing a basis for future closed-loop and neurofeedback applications targeting spatial attention.

## Introduction

Accessing information generated by the brain in real time is a major challenge in systems and cognitive neuroscience. Reliable decoding of brain activity has important applications for brain–computer interfaces, neuroprostheses, and neurofeedback, while also providing a means to investigate the neural representations underlying cognition. In particular, advances in multivariate analysis and machine-learning approaches have made it increasingly possible to extract behaviorally relevant information from distributed patterns of brain activity. Real-time fMRI provides an attractive framework in this context because it combines whole-brain coverage with relatively high spatial resolution and can be used to target specific patterns of activity at the individual-participant level (Taschereau-Dumouchel et al., 2022). Here, we focus on real-time, spatially resolved access to a particularly elusive cognitive state: the locus of covert spatial attention, or attentional spotlight.

Visual spatial attention enables behaviorally relevant information to be prioritized at selected locations in the visual field. It can operate overtly, through movements of the eyes towards relevant stimuli, or covertly, by shifting attention while maintaining fixation (Posner, 1980). Covert spatial attention enhances perceptual sensitivity and behavioral performance at attended locations (Carrasco and Yeshurun, 2009) and recruits a distributed network involving frontoparietal areas, including the frontal eye fields (FEF) and intraparietal sulcus (IPS), together with modulation of sensory representations in visual cortex (Boshra and Kastner, 2022; Corbetta et al., 2008a; Corbetta and Shulman, 2002a). Importantly, spatial attention is not necessarily maintained continuously at a single location. Behavioral and electrophysiological evidence indicates that attentional selection exhibits substantial temporal dynamics and can rhythmically sample competing locations (Fiebelkorn et al., 2013; Fries, 2023; Gaillard et al., 2020; Landau and Fries, 2012). Recent non-invasive MEG decoding further supports this view, showing task-dependent fluctuations in the decoded attentional locus within the alpha frequency range and a relationship between decoding strength and behavioural performance (Mostafalu et al., 2026). Thus, obtaining an instantaneous estimate of covert attention requires capturing a spatially specific but intrinsically dynamic neural state.

Considerable progress has been made towards this objective. In non-human primates, intracortical recordings have enabled high-resolution decoding of the spatial locus of covert attention, including continuous estimates of its position in the visual field (Astrand et al., 2020, 2016; De Sousa et al., 2021; Gaillard et al., 2020). These approaches have demonstrated that trial-by-trial fluctuations in decoded attention are related to perceptual performance and have revealed the fine temporal dynamics of attentional exploration. Non-invasive electrophysiological approaches in humans have also progressed substantially. In particular, covert shifts of attention towards one of four spatial targets can be decoded from single-trial EEG signals (Reichert et al., 2020), demonstrating that single-trial decoding is not restricted to invasive recordings. More recently, MEG-based machine-learning approaches have provided time-resolved estimates of covert spatial attention and revealed rhythmic fluctuations in these decoded representations (Mostafalu et al., 2026). EEG and MEG therefore offer excellent temporal resolution, but obtaining fine-grained spatial information about the locus of attention remains more challenging.

fMRI provides complementary information because attentional modulation can be resolved within the retinotopic organization of the visual cortex. Earlier studies demonstrated that attended spatial locations or visual-field quadrants could be discriminated from multivariate BOLD signals (Andersson et al., 2012; Ekanayake et al., 2018). Recent work has considerably strengthened this observation. (Meyyappan et al., 2025) showed that anticipatory spatial-attention states can be decoded from multivoxel activity throughout the visual cortical hierarchy, including early visual cortex, and that decoding accuracy predicts subsequent behavioral performance. Importantly, these effects were replicated when the attended location was externally instructed or internally chosen, supporting the interpretation that multivariate BOLD patterns contain information about top-down attentional states rather than merely reflecting external cue information. Even more directly, (Bloem et al., 2025) demonstrated that the spatial locus and extent of covert attention can be reconstructed from retinotopically organised BOLD activity in early visual cortex. Their results establish that fMRI can provide fine-grained estimates of the spatial distribution of covert attention under controlled fixation. A critical issue in such decoding approaches is the distinction between covert attention and oculomotor activity. Covert and overt orienting recruit partly overlapping neural systems, although their multivariate representations can be dissociated within the frontoparietal network (Wu et al., 2022). Moreover, covert attention is associated with systematic changes in fixational eye movements, including microsaccades. These signals covary with neural signatures of attention, although attention-related neural modulation can occur in their absence (Liu et al., 2022). Eye position therefore needs to be carefully controlled when interpreting spatially resolved decoding of covert attention. Similarly, strongly salient spatial stimuli may produce location-specific sensory activity that is difficult to distinguish from attentional modulation. Paradigms aimed at isolating covert attention should therefore minimise stimulus-driven spatial information while ensuring stable fixation.

Together, these findings demonstrate that covert spatial attention can be decoded non-invasively and that fMRI can recover fine-grained spatial information about the attentional field. However, an important challenge for real-time applications is to obtain reliable, participant-specific estimates of the attended spatial location on individual trials using a pipeline that can operate online. Such an approach should additionally establish that the decoded information is reproducible within individuals, reflects attentional rather than oculomotor or stimulus-driven signals, and is meaningfully related to behavioral performance.

In the present study, we developed a participant-specific fMRI preprocessing and decoding pipeline designed to estimate the locus of covert spatial attention in real time. Participants performed a cued target-detection task using low-salience visual stimuli, thereby maximising the contribution of endogenous spatial-attention processes while minimising stimulus-driven spatial information. We show that the attended location can be reliably decoded at the single-trial level and that decoding performance remains stable across sessions acquired several days apart. Decoding errors are spatially structured, occurring preferentially at locations neighbouring the cued position rather than being randomly distributed across the visual field. The voxels contributing to decoding follow the retinotopic organisation of occipital visual cortex, consistent with spatially specific attentional modulation of visual representations.

Taken together, our results demonstrate that spatially specific information about covert attention can be extracted from non-invasive fMRI signals at the single-trial and individual-participant levels using a pipeline designed for real-time implementation. By combining spatially resolved decoding, low-salience stimulation, control of eye position, and cross-session reproducibility, this approach provides a framework for tracking covert attentional states in real-time and for the development of future closed-loop and neurofeedback applications targeting spatial attention.

## Materials and Methods

### Participants

Thirteen participants (9 females, mean age: 23.78 years, standard deviation: 5.29 years; 4 males, mean age: 29.25 years, standard deviation: 7.18 years) participated in the experiment after signing written informed consent. All subjects were healthy with no history of psychiatric or neurological disorders and had normal or corrected-to-normal vision. The experimental protocol was approved by the French Ethic Committee (Comité de protection des personnes, #2018-72, ID RCB: 2017-A03612-51), and complied with the Declaration of Helsinki.

### Experimental setup

Octave and the Psychophysics Toolbox (Brainard, 1997; Kleiner et al., 2007; Pelli, 1997) were used to generate the visual stimuli and control the experimental procedure within the scanner environment. The visual stimuli were back projected on a screen outside of the bore at a viewing distance of 130 cm and subjects were able to see the display through a mirror placed in front of their eyes. Subjects, lying in the scanner, were instructed to maintain their gaze fixated on a black cross located at the center of the screen against a gray background. Monocular right eye movements of the participants were recorded during scanning process with an EyeLink 1000 fMRI-compatible eye tracker (SR-Research) sampling at 1000 Hz. Eye calibration was performed before each functional scan. Trials were interrupted if the eyes moved outside the 2°x2° eye control window. Interrupted trials were canceled and re-played at the end of the block.

### Behavioral task

Participants were asked to perform a spatially cued target-detection task in the MRI scanner (Figure 1A). Each trial began with a central fixation cross (0.5° × 0.5° of visual angle) presented for 1.5–2.5 s, followed by a 200-ms central arrow cue (1° × 1°) indicating one of eight possible target locations. Target locations were arranged at 5° eccentricity along the horizontal vs diagonal locations and were marked by small gaps in a circle centered on fixation. Participants were instructed to maintain central fixation while covertly directing their attention toward the cued location.

**Figure 1.**
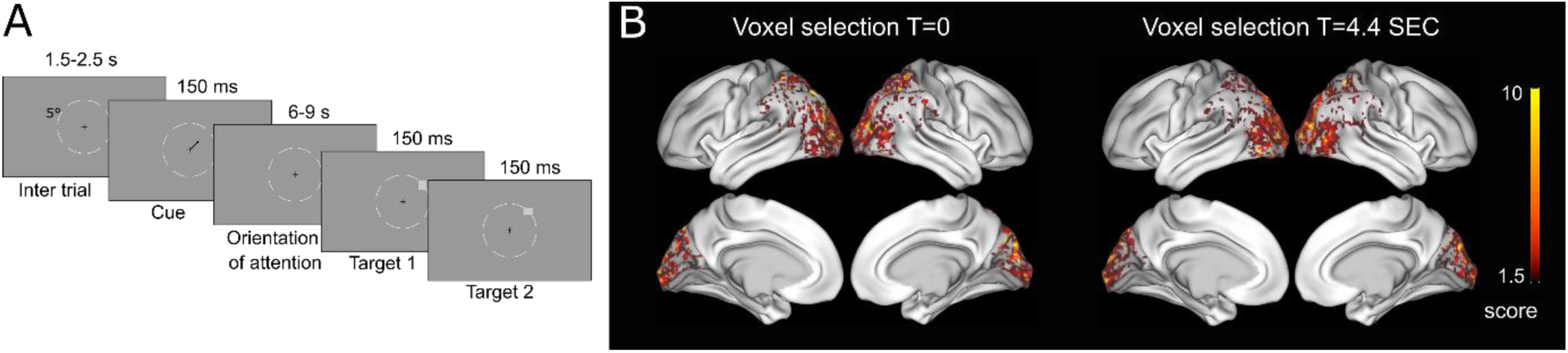
Spatial-attention decoding task and feature selection. (A) Schematic representation of the spatially cued target-detection task. Each trial began with a central fixation period of 1.5–2.5 s, followed by a central arrow cue indicating one of eight possible target locations positioned at 5° eccentricity. Participants maintained central fixation while covertly directing attention toward the cued location. After a variable cue-to-target interval of 6–9 s, two low-contrast oriented targets were presented consecutively at the cued location, separated by a 150-ms interval. Participants reported whether the two targets had the same or different orientations. (B) Group-level spatial distribution of voxels retained during the feature-selection procedure for decoding. Maps are shown for feature selection at target onset (T = 0 s) and 4.4 s after target onset (T = 4.4 s), illustrating the cortical distribution of informative voxels used for spatial-attention decoding. The color scale indicates the feature-selection F score.

After a variable cue-to-target interval of 6–9 s, two low-contrast gray rectangles were presented consecutively at the cued location for 200 ms each, separated by a 150-ms interval. Targets were oriented either vertically (0.5° × 1°) or horizontally (1° × 0.5°). Cue validity was 100%. Participants indicated whether the two targets had the same or different orientations by pressing one of two buttons on a joystick held in their right hand. Responses were required within 1 s of the onset of the first target.

Every five trials, a 10–13-s fixation baseline was presented during which participants maintained central fixation without directing attention toward any specific target location. Each session comprised four blocks of 24 trials presented in randomized order, with three trials per cued location. Block duration ranged from approximately 5 to 10 min, depending on the number of canceled trials. Before each scanning session, target size was individually calibrated to yield approximately 70% correct performance using a 60-trial staircase procedure. Each participant completed four decoding sessions, with two sessions per week over two weeks.

### fMRI scanning procedure

Neuroimaging data were acquired with a 3T MRI scanner (Siemens Prisma) using a 64-channel head coil. Subjects were lying on their back with their head restrained using foam head rest. Functional images with Blood-Oxygen Level Dependent (BOLD) contrast enhancement were acquired for each participant. Single-shot echo-planar images (EPIs) acquired using a typical T2*-weighted gradient-echo, echo-planar imaging sequence (TR/TE =2200/30ms; flip angle, 90°; 3mm isotropic voxels, 40 interleaved ascending slices with no gap) with a 74×74 acquisition matrix and a 518×518 mm rectangular field of view. A High-resolution T1 structural image (MPRAGE, 0.875mm3 isotropic voxels, slices, TR/TE=3.5/3.42s, flip angle, 8°) was also acquired for each single subject, during the first fMRI session.

### Preprocessing for standard event-related analyses

Image pre-processing was performed using Statistical Parametric Mapping (SPM12) (Friston, 1994) software (Wellcome Centre for Human Neuroimaging, University College London,UK; 204 http://www.fil.ion.ucl.ac.uk/spm). To adjust within-volume time differences, Temporal misalignment between different slices of the functional data, introduced by the sequential nature of the fMRI acquisition protocol, is corrected using SPM12 slice-timing correction (STC) procedure (Henson et al. 1999), where the functional data is time-shifted and resampled using sinc-interpolation to match the time in the middle of each TA (acquisition time). Functional data were then realigned using SPM12 realign & unwarp procedure (Andersson et al., 2001), where all scans are resampled to a reference image (first scan of the first session) using b-spline interpolation. This procedure also addresses potential susceptibility distortion-by-motion interactions by estimating the derivatives of the deformation field with respect to head movement, and resampling the functional data to match the deformation field of the reference image. Functional scans were then co-registered with the 3D structural image and spatially normalized against the standard, stereotactic space of the MNI template. Spatial smoothing was performed with a Gaussian kernel of 8-mm full width at half maximum. A single subject, fixed-effect model was created for each individual participant in order to perform the based conditions random-effect analysis. The motion parameters were included in the General Linear Model (GLM) as regressors to correct the head movement.

### SPM Contrasts

In order to describe activations elicited by the decoding task, during attention orientation, two contrasts were used; Attention>Fixation and Right>Left. Within both contrasts attention and right/left conditions were sampled in an interval of 2 seconds before target onset. all trials were pooled together, irrespective of the cued position within the trial.

### Decoding pre-processing procedure

For the decoding, and all the following analyses, the functional images were processed in real time using the FRIEND toolbox (the institut d’Or, Brazil) (Basilio et al., 2015; Sato et al., 2013). This allowed a simplified version of preprocessing pipeline. During the acquisition the functional images were transformed in the NIFTI format with the dicom2nii toolbox (Li et al., 2016) and realigned to one reference anatomical image using FSL tools (Jenkinson et al., 2012). No smoothing was implemented as we wanted to keep each vowel as an independent feature. Voxel activity was extracted for each trial and normalized by its mean activity recorded during the fixation period preceding the trial. Prior to decoding, voxels of interest were chosen based on an ANOVA, testing the significance of the activity of each voxel to one or several of the cued locations during the cue to target interval. Only the significant voxels (p<0.05) were selected for decoding using the python neuroimaging toolbox Nilearn (Abraham et al., 2014).

### Decoding procedure

The decoding procedure was performed offline. Following preprocessing, informative were selected using an ANOVA an only those showing a significant effect (p < 0.05) were retained as features for subsequent decoding. Then, a linear support vector machine (SVM) with a one-vs-all classification scheme was then trained to associate multivoxel BOLD activity patterns with the corresponding spatial cueing condition. The decoder was designed to classify each activity pattern into one of the eight possible spatial locations. Training data consisted of correct trials acquired during the four decoding sessions.

Decoding performance was assessed using a leave-one-block-out cross-validation procedure. For each fold, voxel selection and classifier training were performed using data from all blocks except one. Training trials were balanced across spatial locations to ensure that each of the eight classes contributed an equal number of trials. The trained classifier was then tested on the held-out block, which therefore contained only trials that had not been used for either voxel selection or classifier training. This procedure was repeated until each block had served once as the independent test set, and decoding accuracy was averaged across folds to obtain an overall measure of classifier performance.

For the temporal analysis of decoding performance, this cross-validation procedure was repeated independently for successive fMRI volumes, from 2.2 s before cue onset to 6.6 s after target onset, allowing the time course of spatial-attention decoding to be assessed. Voxel selection was performed using Nilearn (Abraham et al., 2014), whereas SVM classification and decoding analyses were implemented using MATLAB’s Statistics and Machine Learning Toolbox.

### Statistical analyses

Statistical analyses were performed at the group level across participants. Unless otherwise specified, group data are reported as mean ± SEM, and statistical significance was assessed using two-tailed tests with an alpha level of 0.05. For the eight-class decoding analyses, theoretical chance performance was 12.5%. Decoding performance relative to chance was assessed using permutation-based estimates of the null distribution, from which a 95% confidence interval around chance was derived. Decoding performance was considered above chance when it exceeded the upper bound of this permutation-derived confidence interval.

Because several analyses involved repeated measurements obtained from the same participants and the assumptions required for parametric paired tests were not assumed, paired comparisons were performed using two-sided Wilcoxon signed-rank tests. These tests were used to compare decoding performance between upper-and lower-visual-field locations and between diagonal and horizontal locations at each angular offset, as well as to compare the proportion of ipsilateral and contralateral attention-selective voxels within occipital and parietal cortices. Associations between decoding performance and the proportion of attention-related voxels across participants were assessed using Spearman rank correlations (Spearman’s ρ). This non-parametric correlation was used to quantify monotonic relationships without assuming normally distributed variables.

For conventional SPM analyses, whole-brain group-level activation maps were thresholded at p < 0.05, family-wise error (FWE) corrected.

## Results

In this study, our goal was to assess the feasibility of real-time decoding of spatial attention from fMRI BOLD signal and characterize the stability of this decoding in time, within and across sessions. We thus had subjects perform a spatial attention centrally cued target detection task while their brain activity was scanned using a 3T MRI scanner. Attention could be cued towards one out of 8 possible spatial position (Figure 1A). Subjects sat in four scanning sessions. In each session they performed 5 blocks of 24 trials per block. In the following, we describe the whole brain activity associated with spatial attention then we focus on the single trial decoding of spatial attention orientation.

### Whole-brain activity associated with spatial attention

Whole-brain analyses reproduced prior observations (Corbetta et al., 2008b; Corbetta and Shulman, 2002b; Simpson et al., 2011) and revealed a distributed pattern of activity associated with the attentional task (supplementary Figure S1). The Attention > Fixation contrast showed robust bilateral activation across occipital, parietal, and frontal regions. Significant clusters were observed in the middle occipital cortex, superior and inferior parietal cortices, middle frontal and precentral regions, supplementary motor area, postcentral cortex, and insular cortex. Overall, this activation pattern is consistent with the recruitment of a distributed frontoparietal and occipital network during spatial attention.

To assess the spatial selectivity of these responses, we further contrasted trials in which attention was directed toward the right versus the left visual field. The Right > Left contrast revealed a predominantly left-lateralized pattern of activity within occipital and ventral occipitotemporal cortex, including the left middle and inferior occipital cortices, fusiform cortex, and lingual gyrus. This contralateral organization indicates that directing attention toward the right visual field was associated with stronger responses in left visual cortical regions compared with directing attention toward the left visual field. Together, these results show that the task engaged the expected large-scale attention-related network while also producing spatially specific, contralateral modulations within visual cortex.

### Spatial distribution of attention-discriminative voxels: visual and parietal cortex

To characterize the spatial distribution of the features contributing to the decoding analysis, we examined the voxels identified by the ANOVA as being modulated by the cued attentional location. Feature selection was performed independently for each participant, and the resulting subject-specific maps were averaged across participants (Figure 1B). Thus, the group maps shown here provide a spatial summary of the regions consistently selected across participants.

At target onset (T = 0 s), selected voxels were predominantly distributed bilaterally across posterior cortical regions, with a strong concentration in the occipital cortex and additional contributions extending into dorsal posterior/parietal regions. This spatial distribution indicates that information discriminating the eight attended locations was largely represented within visually responsive and spatial-attention-related cortical areas. A qualitatively similar distribution was observed 4.4 s after target onset (T = +4.4 s). At this later time point, informative voxels remained strongly concentrated within bilateral occipital regions, with additional posterior dorsal contributions. The substantial overlap between the two time points suggests that spatially informative patterns were supported by a relatively consistent set of posterior cortical regions across the attentional and early post-target periods. Nevertheless, the stronger and more extensive occipital contribution at T = +4.4 s is also compatible with the additional contribution of target-evoked visual activity at this time point.

Overall, these group-level feature-selection maps show that the voxels carrying information about the locus of covert attention are primarily located within visual cortex, together with additional parietal contributions. The predominance of occipital features is consistent with the regional decoding analyses reported below, showing that occipital voxels account for a substantial proportion of the spatial information available to the decoder.

### Spatio-temporal dynamics of covert spatial attention decoding

Based on this subject-specific feature selection, we first assessed whether the spatial locus of covert attention could be decoded from single-trial fMRI activity and examined how decoding performance evolved over time. For each participant, a linear support vector machine was trained to discriminate among the eight possible attended locations using a leave-one-block-out cross-validation procedure. Feature selection was performed independently within each training fold to identify voxels whose activity was modulated by the cued attentional location. Figure 2A shows decoding performance as a function of time relative to target onset for each participant (n = 13) and at the group level. Given the eight possible spatial locations, theoretical chance performance was 12.5%. During the cue period, decoding remained close to chance, with a mean accuracy of approximately 12–13%. Performance subsequently increased, reaching approximately 19% at −4.4 s and 30% around target onset and remained at a comparable level (32%) at +2.2 s.

**Figure 2.**
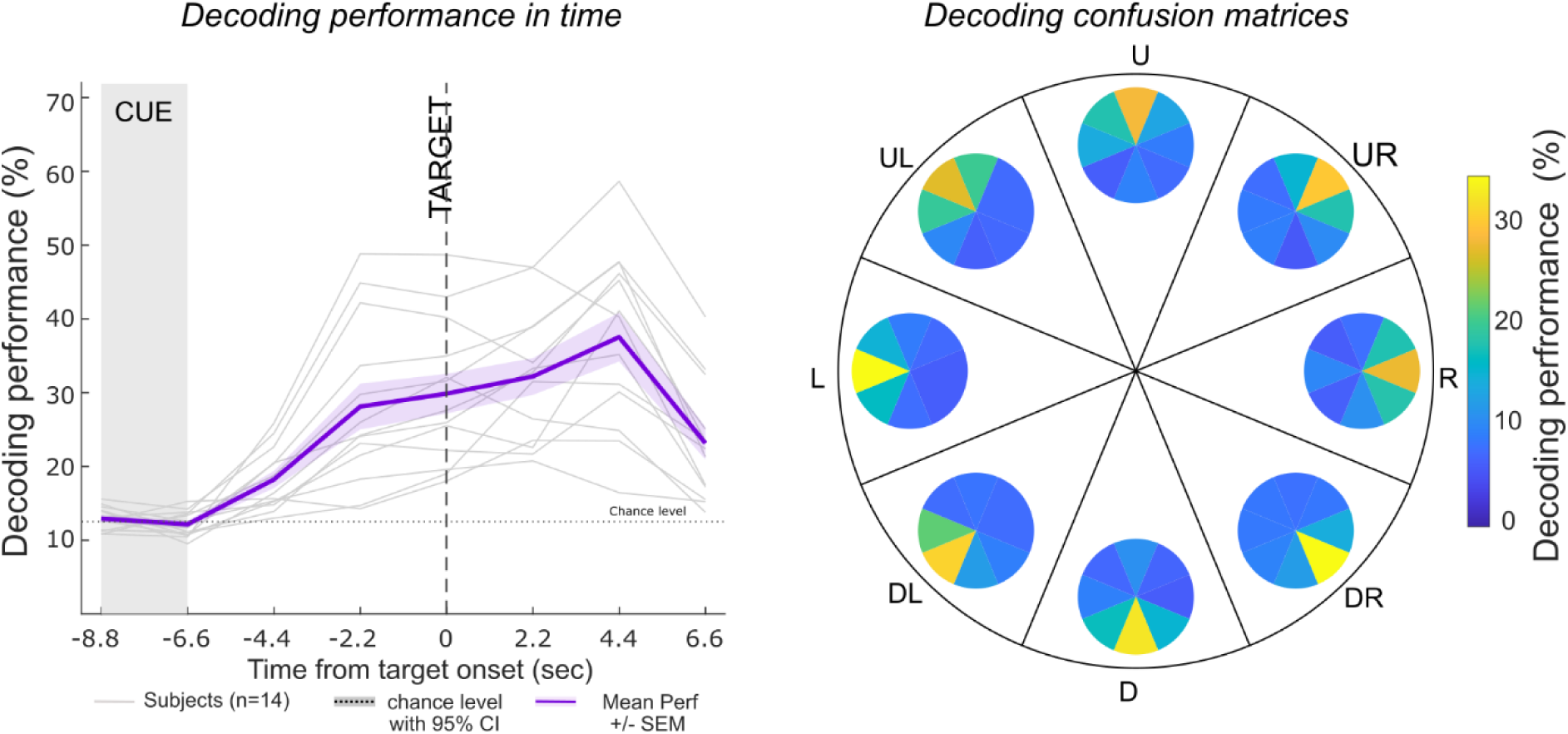
Temporal dynamics and spatial organization of covert-attention decoding. (A) Decoding performance for the eight possible covert-attention locations as a function of time relative to target onset. Thin gray lines represent individual participants (n = 13), and the purple line represents the group mean ± SEM. The horizontal dashed line indicates the theoretical chance level (12.5%), with the permutation-derived 95% confidence interval shown around chance. The shaded area indicates the cue period, and target onset is indicated by the vertical dashed line at 0 s. (B) Spatially organised decoding confusion matrices for the eight possible attentional locations (U, UR, R, DR, D, DL, L, and UL). Colour indicates decoding performance (%).

Group-average decoding performance reached a maximum of approximately 38% at +4.4 s after target onset, corresponding to more than three times the theoretical chance level. At this time point, decoding accuracy reached approximately 60% in the best-performing individual participants. Performance subsequently decreased to approximately 23–24% at +6.6 s. At the group level, decoding performance rose above the permutation-derived 95% confidence interval around chance from approximately −4.4 s relative to target onset and remained above this level throughout the subsequent time points. The temporal evolution of decoding performance is consistent with the expected hemodynamic delay of the BOLD response. The additional increase observed approximately 4.4 s after target presentation may reflect the contribution of target-evoked activity superimposed on the ongoing attentional representation. Together, these results demonstrate that the spatial locus of covert attention can be reliably decoded at the single-trial level across eight possible spatial locations.

### Spatial organization of decoding errors

We next examined whether classification errors were randomly distributed across spatial locations or instead retained information about the position of the attentional locus. To this end, we computed group-level confusion matrices for the eight possible cued locations using decoding estimates obtained at target onset (T = 0 s; Figure 2B). This time point was selected because decoding performance was already substantially above chance while preceding the delayed BOLD response associated with target presentation. For each attentional condition, the highest decoding probability was associated with the correctly cued location. Importantly, however, incorrect classifications were not uniformly distributed across the seven remaining positions. Instead, decoding errors occurred preferentially at spatial locations neighbouring the cued position, whereas more distant locations were less frequently confused with the attended location.

This spatial organization indicates that the decoder preserved information about the topography of covert spatial attention even when the exact cued location was not correctly classified. In other words, classification errors were predominantly local rather than arbitrary: when the decoder failed to recover the exact attentional locus, its prediction generally remained spatially close to the true attended position. The structured pattern of decoding errors therefore suggests that the classifier captured a graded spatial representation of the attentional locus rather than simply discriminating eight unrelated experimental conditions. Such neighbouring-location errors may arise partly from noise in the BOLD signal, but they are also compatible with trial-to-trial variability in the precise spatial deployment of covert attention. This interpretation is consistent with electrophysiological evidence indicating that the attentional spotlight is intrinsically dynamic rather than continuously fixed at a single spatial position (Fries, 2023; Gaillard et al., 2020).

Together, the temporal decoding profile and the spatial structure of classification errors demonstrate that single-trial fMRI activity contains spatially specific information about the locus of covert attention across eight possible locations.

### Contribution of parietal and occipital voxels to attention decoding

To determine the relative contribution of occipital and parietal cortices to the decoding of covert spatial attention, we compared decoding performance obtained from four sets of informative voxels: occipital voxels, parietal voxels, combined occipital–parietal voxels, and all informative voxels identified at the whole-brain level. Occipital and parietal regions of interest (ROIs) were defined using anatomical masks derived from the Harvard–Oxford Cortical Structural Atlas (Desikan et al., 2006; Frazier et al., 2005; Goldstein et al., 2007; Makris et al., 2006)and mapped to each participant’s functional space.

### Temporal decoding profiles across cortical regions

Decoding performance differed markedly depending on the anatomical origin of the selected voxels (Figure 3A). Whole-brain, occipital, and combined parietal–occipital decoding showed highly similar temporal profiles. Performance was close to the theoretical chance level of 12.5% during the early cue period, increased to approximately 19% at −4.4 s relative to target onset, and reached approximately 28% at −2.2 s. At target onset, decoding accuracy was approximately 30% and slightly higher at +2.2 s (32%). Performance subsequently peaked at +4.4 s, reaching approximately 37–38% for the whole-brain decoder and approximately 34–35% for the occipital and combined parietal–occipital decoders, before decreasing to approximately 23–24% at +6.6 s. In contrast, decoding based exclusively on parietal voxels remained close to chance throughout the trial, with performance fluctuating around approximately 11–15%. Parietal decoding did not show the marked pre-target increase or the post-target peak observed for the other voxel sets.

**Figure 3.**
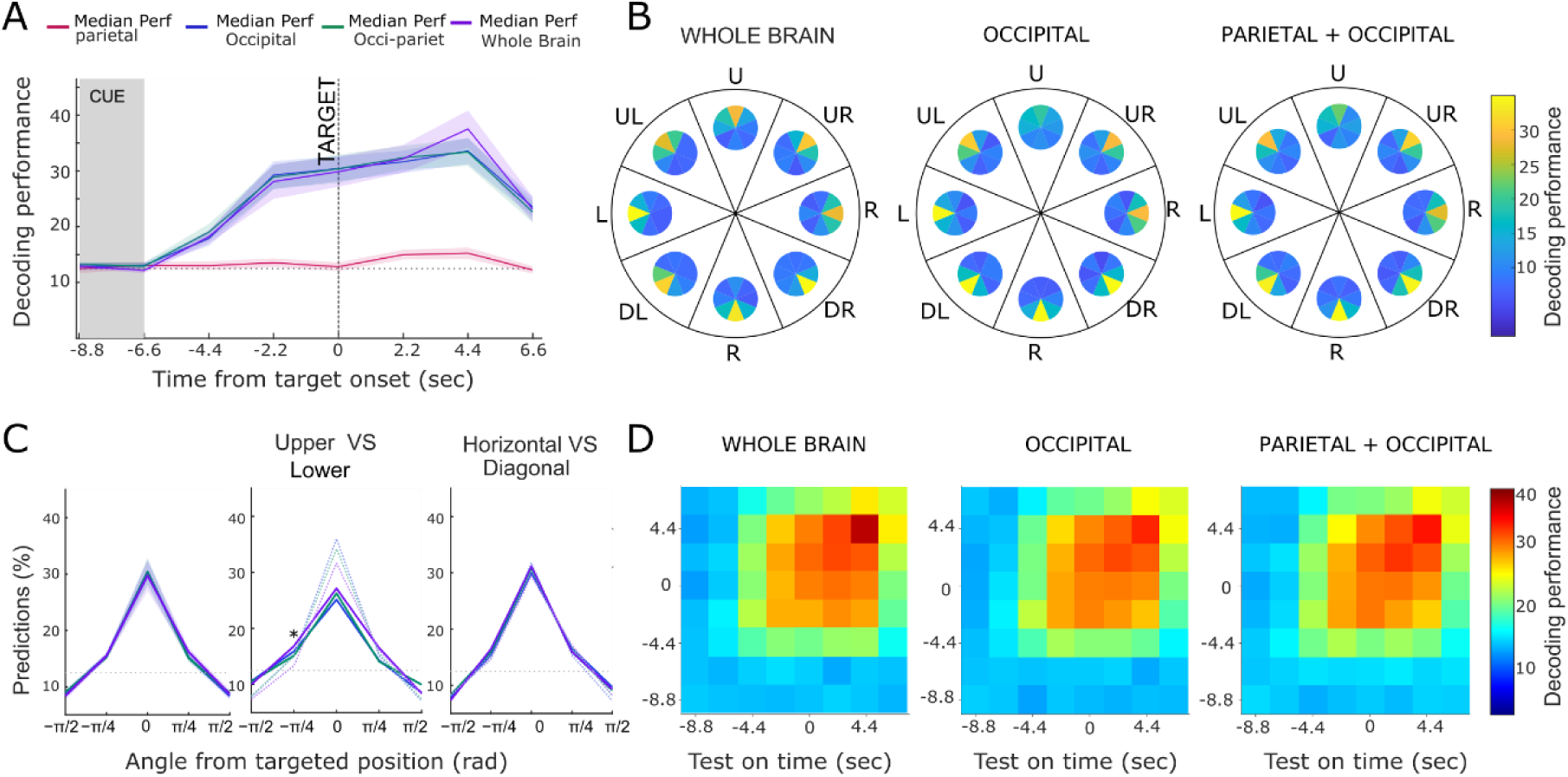
Regional contributions to the spatial and temporal decoding of covert attention. (A) Time-resolved decoding performance obtained using informative voxels selected from the parietal cortex, occipital cortex, combined parietal and occipital cortices, or the whole brain. Decoding performance is shown as a function of time relative to target onset. The shaded region indicates the cue period and the vertical dashed line indicates target onset. The horizontal dotted line represents the theoretical chance level for eight-class decoding (12.5%). Lines represent group-level decoding performance and shaded areas indicate ± SEM. (B) Spatial confusion matrices at target onset (T = 0 s) for whole-brain, occipital, and combined parietal– occipital decoding. The eight sectors correspond to the eight cued attentional locations (U, UR, R, DR, D, DL, L, and UL), and each internal pie chart represents the distribution of decoder predictions for a given cued location. Colors indicate decoding performance (%). (C) Spatial distribution of decoder predictions expressed as a function of angular distance from the cued position. Left: predictions pooled across all eight attentional locations. Middle: comparison of upper-and lower-visual-field locations). Right: comparison between locations positioned along horizontal vs diagonal locations. (D) Cross-temporal generalization matrices for whole-brain, occipital, and combined parietal–occipital decoding. Classifiers were trained independently at each time point (y-axis) and tested at every other time point (x-axis). Color indicates decoding performance (%).

These results indicate that the fine-grained spatial information required to discriminate among the eight attended locations is predominantly carried by occipital activity. Importantly, restricting the decoder to occipital voxels reproduced most of the temporal profile obtained at the whole-brain level, whereas the addition of parietal voxels provided only limited improvement.

### Spatial organization of regional decoding errors

We next examined whether the spatial organization of classifier predictions depended on the anatomical origin of the contributing voxels (Figure 3B). Confusion matrices were computed at target onset (T = 0 s), when decoding performance was already substantially above chance but preceded the delayed BOLD response to target presentation. Whole-brain, occipital, and combined parietal–occipital decoders displayed very similar spatial confusion patterns. For each cued attentional location, the highest probability was associated with the cued spatial position. Incorrect classifications were not uniformly distributed across the remaining positions but occurred preferentially at locations neighboring the cued position. Conversely, positions located farther from the true attentional locus were less frequently selected by the decoder. Thus, the characteristic topographical organization of decoding errors observed at the whole-brain level was largely preserved when decoding was restricted to occipital cortex. This finding further indicates that occipital activity contains most of the fine-grained spatial information supporting localization of the covert attentional spotlight.

### Angular organization of decoding errors and visual-field anisotropies

To quantify the spatial organization of these errors, decoder predictions were expressed as a function of their angular distance from the cued position (Figure 3C). Across all voxel sets, prediction rates were strongly centered on the true attentional location (Figure 3C, left). Decoding reached approximately 30–33% at an angular distance of 0 rad for the whole-brain, occipital, and combined parietal–occipital decoders. Prediction rates decreased to approximately 15–16% for immediately neighboring positions (±π/4) and to approximately 8–10% for more distant positions (±π/2). This graded spatial profile confirms that decoding errors were predominantly local rather than randomly distributed across the visual field. We then investigated whether this spatial organization differed between upper and lower visual-field locations (Figure 3C, middle). A modest upper–lower asymmetry was observed. Prediction rates at the actual cued position were numerically higher for lower-than upper-field targets, although this difference did not reach statistical significance in the whole-brain decoder (p = 0.09). The only significant upper–lower difference was observed at the −π/4 angular offset (p = 0.04), where prediction rates were higher for upper-than for lower-field targets. No significant upper–lower differences were observed at the remaining angular offsets (all p ≥ 0.3). Thus, the upper–lower asymmetry was spatially specific rather than reflecting a consistent overall advantage for either visual field. Such an anisotropy is consistent with previously reported behavioral and neurophysiological asymmetries in visual spatial processing (Carrasco et al., 2004; Ibos et al., 2009; Montaser-Kouhsari and Carrasco, 2009; Zénon et al., 2009). In contrast, separating locations according to whether they were positioned along the horizontal versus diagonal locations revealed highly similar spatial decoding profiles (Figure 3C, right). In both configurations, prediction rates peaked at approximately 32–35% for the correct location and decreased similarly with increasing angular distance from the target. No reliable difference between horizontal and diagonal locations was observed at any angular offset (all p ≥ 0.1). Thus, the only reliable spatial anisotropy identified by the decoder concerned the upper versus lower visual field at the −π/4 angular offset, rather than the distinction between horizontal and diagonal locations.

### Within trial decoding stability

The analyses described above characterize decoding performance independently at each time point. We next asked whether the spatial information captured by the decoder relied on a stable neural representation across time or, alternatively, on temporally specific patterns that changed throughout the trial. To address this question, we performed a cross-temporal generalization analysis in which classifiers trained at each time point were tested on activity patterns acquired at all other time points (Figure 3D). At the whole-brain level, decoding generalized across a broad temporal window surrounding target onset. Classifiers trained during the late cue-to-target interval successfully decoded the attended location at neighboring time points, producing an extended region of above-chance performance around the train–test diagonal. Cross-temporal generalization emerged from approximately −4.4 s relative to target onset and extended throughout the period surrounding target presentation. The strongest decoding was observed when training and testing were performed during the later part of this interval, particularly around +4.4 s, where performance reached approximately 38%.

A highly similar cross-temporal structure was observed when decoding was restricted to occipital voxels or to the combined occipito-parietal voxel set. In both cases, classifiers trained during the period surrounding target onset generalized across several neighboring time points, and the overall temporal organization closely resembled that obtained using whole-brain informative voxels. The strongest cross-temporal decoding was again observed during the late peri-target and early post-target periods. Importantly, generalization was not restricted to identical training and testing time points. Instead, the broad off-diagonal generalization indicates that the spatial code underlying the decoded attentional location remained sufficiently similar across successive BOLD samples to support decoding across time. At the same time, decoding performance varied across the trial time, indicating a progressive strengthening of the spatial representation rather than a temporally invariant signal.

These results therefore suggest that the neural representation of the attended spatial location is relatively stable over the cue-to-target and early post-target periods. Moreover, the close similarity between the whole-brain, occipital, and occipito-parietal cross-temporal generalization patterns indicates that this temporally stable spatial representation can largely be recovered from occipital activity. Together with the regional decoding analyses, these findings further support a predominant contribution of occipital cortex to the fine-grained representation of the covert attentional locus.

### Within trial attentional spotlight location variability

Although the cross-temporal analyses indicate that the neural representation of the attended location is stable at the population level across successive time points, this does not necessarily imply that the decoded attentional locus remains stationary within individual trials. Previous behavioral and electrophysiological studies have shown that spatial attention can fluctuate over time(Gaillard et al., 2020; Gaillard and Ben Hamed, 2022). We therefore examined the within-trial stability of the decoded attentional location during the period preceding target presentation. For this analysis, we selected trials for which the decoder correctly identified the cued attentional location at target onset (T = 0 s). We then examined the decoder predictions for these same trials at 2.2 and 4.4 s before target onset and expressed them according to their angular distance from the cued location (Supplementary Figure S2). A value of 0 rad therefore corresponds to trials for which the attentional location was already correctly decoded at the earlier time point, whereas increasing angular distances indicate progressively larger spatial deviations from the cued position. The distribution of decoder predictions became more concentrated around the cued location as target onset approached. At −2.2 s, 36.4% of trials that were correctly classified at T = 0 s were already classified at the cued location, compared with 22.3% at −4.4 s. At both time points, incorrect predictions were distributed across the remaining angular positions, with a tendency for errors to occur more frequently at locations relatively close to the cued position. Thus, trials that ultimately yielded a correct estimate of the attentional locus at target onset did not necessarily exhibit the same decoded spatial position throughout the preceding cue-to-target interval. Instead, the probability of recovering the correct location increased as target presentation approached. This temporal evolution is consistent with a progressive stabilization of the decoded attentional locus around the task-relevant location, while also revealing substantial trial-to-trial variability in its estimated spatial position.

Importantly, these results complement rather than contradict the cross-temporal generalization analysis. While cross-temporal decoding demonstrates that a common spatial code can generalize across successive BOLD samples at the population level, the present analysis reveals variability in the location assigned by the decoder on individual trials. Together, these findings suggest that the neural code supporting spatial attention is sufficiently stable to be decoded across time, while the trial-wise decoded estimate of the attentional locus remains temporally variable.

### Cortical topography of attentional processes

To further characterize the relationship between the decoder and the functional organization of visual and attentional cortical systems, we identified, for each attention-selective voxel, its preferred attentional position and projected this information onto cortical flat maps. Figure 4A illustrates the resulting spatial organization of attentional selectivity. In occipital cortex, attention-selective voxels exhibited a clear and orderly spatial pattern. Voxels preferring different attended locations were distributed across posterior visual areas in a manner consistent with the known retinotopic organization of occipital cortex. In particular, responses associated with different sectors of the visual field formed distinct spatially organized clusters, indicating that the information used by the decoder preserved the topographical layout of visual space. By contrast, selective voxels in parietal cortex showed a coarser spatial organization. Although attention-related voxels were present in posterior parietal regions, their distribution did not reveal the same fine-grained topographic arrangement observed in occipital cortex for the eight attended locations. Rather, parietal selectivity appeared to reflect a broader spatial coding scheme, more readily captured at the level of large-scale spatial categories such as visual hemifields. To quantify the inter-hemispheric organization of attention-selective voxels at the group level, we compared the proportion of unique decoding voxels located in the hemisphere contralateral versus ipsilateral to the attended hemifield within occipital and parietal cortices (Figure 4B). In the occipital cortex, the proportion of selective voxels was significantly greater in the contralateral than in the ipsilateral hemisphere (p = 0.0002). A similar contralateral predominance was observed in the parietal cortex, with a significantly greater proportion of selective voxels in the contralateral than in the ipsilateral hemisphere (p = 0.001). These results demonstrate a robust contralateral organization of spatial-attention representations in both occipital and parietal cortices.

**Figure 4.**
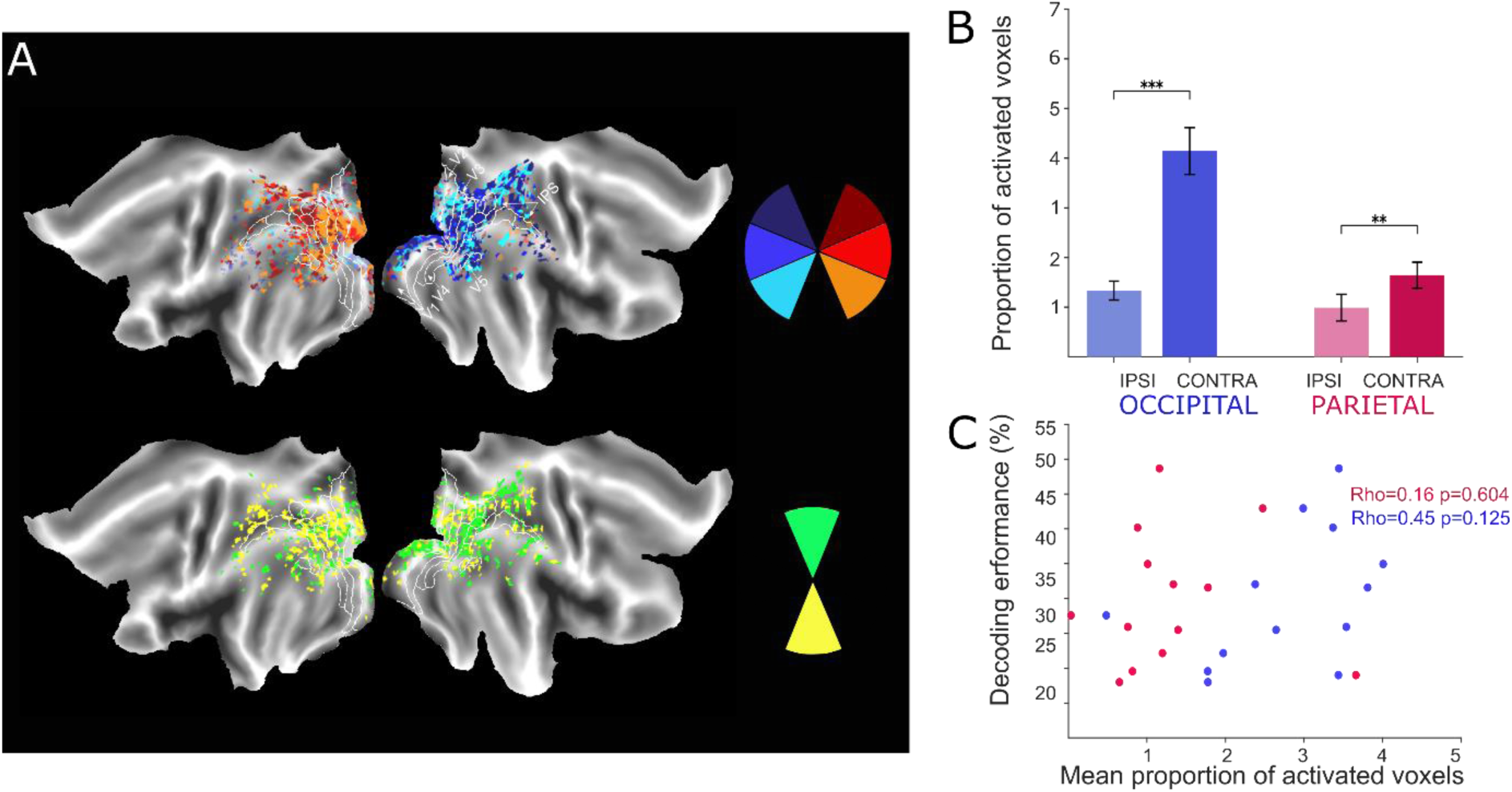
Cortical topography of spatial-attention-related activity. (A) Cortical flat maps showing the spatial distribution of attention-selective voxels for a representative participant. The upper maps illustrate position-selective responses across right and left positions, revealing a fine-grained spatial organization predominantly within occipital cortex. The lower maps show selectivity for up and down positions. White contours indicate the boundaries of the occipital regions, including retinotopic visual areas and parietal regions. (B) Group-level proportion of attention-selective voxels located ipsilateral or contralateral to the attended hemifield. Error bars indicate [SEM/SD]. (C) Relationship across participants between decoding performance and the mean proportion of attention-selective voxels. Each dot represents one participant.

We next examined whether decoding performance covaried across participants with the proportion of attention-selective voxels identified in each region (Figure 4C). No significant correlation was observed between decoding performance and the mean proportion of activated voxels in parietal cortex. A positive but non-significant association was observed in occipital cortex. Thus, although occipital cortex contained a larger contralateral pool of selective voxels and showed a clearer topographic organization, inter-individual variability in the amount of selected voxels did not significantly predict decoding performance in the present sample.

Taken together, these findings show that covert spatial attention is associated with a topographically organized modulation of BOLD activity in visual cortex, consistent with the retinotopic structure of occipital representations. They further indicate that parietal cortex contributes to attentional processing primarily at larger, contralateral spatial scale, whereas the fine-grained location-specific information exploited by the decoder is predominantly expressed in occipital cortex. Most importantly, these results demonstrate that such topographically organized attentional information can be recovered at the single-trial level from fMRI signals.

To further investigate the spatial organization shown in Figure 4, we independently characterized the spatial organization of attention-related BOLD responses using conventional univariate SPM analyses (Supplementary Figure S3). For each of the eight cued positions, activity during spatial attention was contrasted against the fixation baseline, and only voxels showing a significant response were retained (p < 0.05, FWE corrected). This procedure yielded separate activation maps for each attended position, allowing us to examine whether the spatial organization observed among decoder-selected voxels was also apparent in independently defined task-activated voxels.

The resulting maps revealed a spatially structured pattern of activation in occipital cortex, with different attended positions engaging partially distinct portions of visual cortex. In contrast, activations in parietal cortex were more spatially overlapping and showed a less fine-grained organization. Consistent with this pattern, the proportion of activated voxels was significantly greater in the hemisphere contralateral than ipsilateral to the attended hemifield in occipital cortex (p=0.007), whereas no significant contralateral–ipsilateral difference was observed in parietal cortex.

Finally, the mean proportion of activated voxels did not significantly correlate with decoding performance across participants in either occipital or parietal regions. Thus, these independent SPM-based analyses broadly support the spatial organization observed for decoder-selected voxels in Figure 4, particularly the stronger contralateral and position-specific organization in occipital cortex, while also indicating that the overall extent of task-evoked activation alone does not account for individual differences in decoding performance.

### Decoding stability across scanning blocks and sessions

For potential real-time and neurofeedback applications, an important requirement is that decoding performance remains reliable when the decoder is trained on data acquired across different blocks and sessions. We therefore first assessed how decoding accuracy varied as a function of the amount of training data by progressively increasing the number of blocks used to train the classifier and testing it on independent remaining blocks (Figure 5A). Decoding performance was evaluated at target onset (T = 0 s), when spatial-attention information was robustly represented. At the group level, decoding accuracy increased progressively with the number of training blocks. Mean performance was approximately 14% when the decoder was trained on two blocks and increased rapidly as additional training data were included, reaching approximately 25% after 6-8 blocks. Performance continued to improve more gradually thereafter, reaching approximately 26-28% after about 10-12 training blocks. Beyond this point, adding further training blocks produced relatively limited additional improvement, and group-average performance fluctuated around this level. Individual participants showed substantial variability in both decoding performance and learning curves. While some participants reached decoding accuracies of approximately 50-60% with increasing amounts of training data, others stabilized at lower performance levels. Nevertheless, the group-average decoding remained clearly above the theoretical chance level of 12.5% across most training-set sizes once a sufficient number of blocks had been included. We next examined whether decoding performance remained stable across separate scanning sessions (Figure 5B). Mean decoding accuracy was approximately 26% in session 1, 32% in session 2, 29% in session 3, and 31% in session 4. Thus, decoding performance remained substantially above chance across all four sessions, with only moderate variability between sessions and no clear progressive loss of decoding accuracy over time.

**Figure 5.**
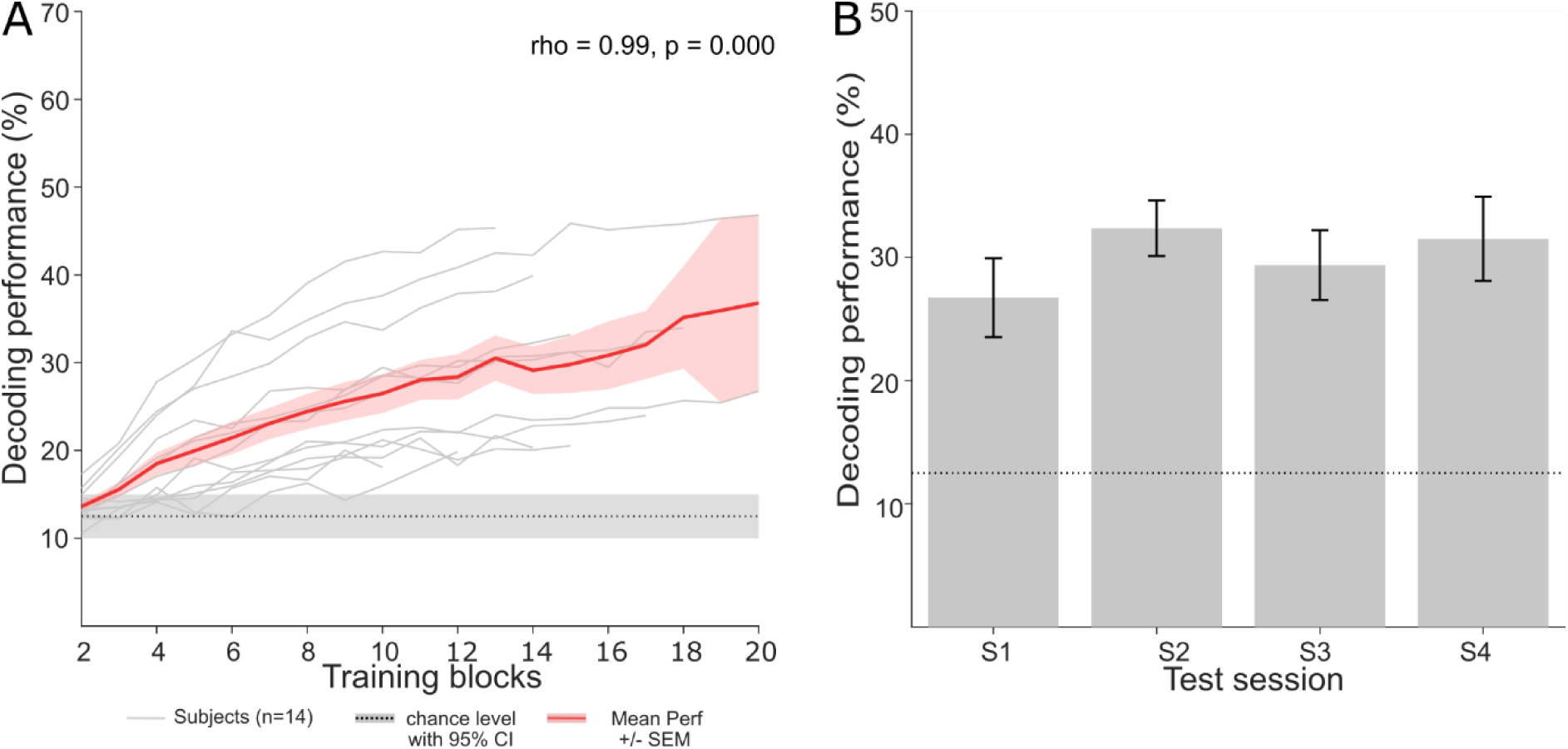
Stability of decoding performance across training blocks and scanning sessions. (A) Decoding performance as a function of the number of training blocks. Thin white lines represent individual participants (n = 13), and the red line shows group mean decoding performance ± SEM. The horizontal dotted line indicates the theoretical chance level for eight-class decoding (12.5%), and the shaded band represents the permutation-derived 95% confidence interval around chance. (B) Mean decoding performance across four independent scanning sessions (S1–S4). Error bars indicate ± SEM.

Together, these results indicate that reliable decoding of covert spatial attention can be achieved after accumulating approximately 10-12 training blocks and that performance remains relatively stable across scanning sessions. The limited improvement observed after this amount of training data suggests that decoding performance is not determined solely by the quantity of training data. Instead, the remaining variability may reflect a combination of measurement noise, inter-individual differences in the strength of spatially specific BOLD representations, and trial-to-trial variability in the deployment of covert attention. This interpretation is consistent with the spatially structured classification errors described above, as well as with the within-trial variability of the decoded attentional locus (Supplementary Figure S2). Thus, although increasing the amount of training data improves classifier performance, the upper limit of decoding accuracy is likely constrained by both properties of the fMRI signal and variability in the attentional state itself.

## Discussion

The present study demonstrates that spatially specific information about covert attention can be extracted from non-invasive 3T fMRI BOLD signals at the single-trial and individual-participant levels. Using an eight-location spatial-attention task, decoding performance substantially exceeded the theoretical chance level of 12.5%, reaching approximately 30% at the group level around target presentation and substantially higher values in the best-performing participants. Importantly, decoding errors were spatially structured, with incorrect predictions preferentially falling near the cued location rather than being uniformly distributed across visual space. Fine-grained spatial information was predominantly carried by occipital activity, whereas parietal voxels showed a broader contralateral organization but provided little information for discriminating among the eight individual locations. Finally, the spatial representation available to the decoder generalized across successive BOLD samples, showed systematic within-trial variation, and remained reproducible across scanning sessions. Together, these findings establish that multivoxel fMRI activity contains sufficiently detailed and reproducible information to estimate the locus of covert spatial attention on individual trials.

### Fine-grained single-trial decoding of covert spatial attention

Previous fMRI studies established that the direction of covert attention can be discriminated from multivariate BOLD activity, including classification among visual-field quadrants (Andersson et al., 2012; Ekanayake et al., 2018). More recent work has substantially extended these findings. (Meyyappan et al., 2025) showed that anticipatory spatial-attention states can be decoded throughout the visual cortical hierarchy and that the strength of these representations is related to subsequent behavioral performance. (Bloem et al., 2025) further demonstrated that both the locus and spatial extent of covert attention can be reconstructed from retinotopically organized activity in early visual cortex.

Our findings complement this literature by showing that participant-specific multivoxel patterns acquired at 3T provide sufficient information to discriminate among eight possible attended locations on individual trials. Increasing the number of spatial classes necessarily makes classification more demanding than binary or quadrant-level decoding, because neighboring attentional states are represented by partially overlapping neural populations. Consequently, absolute decoding accuracy should not be directly compared across studies using different numbers of classes, stimuli, acquisition parameters, or classification procedures. Nevertheless, performance reaching approximately three times the theoretical chance level indicates that the BOLD signal contains substantial fine-grained information about the current spatial locus of attention.

An important observation is that classification errors themselves retained spatial information. Incorrect predictions occurred preferentially at locations adjacent to the cued position, whereas spatially distant locations were less frequently confused. Thus, the classifier did not simply distinguish eight unrelated experimental labels. Rather, the geometry of its errors reflected the geometry of the visual field. This graded organization is compatible with spatially overlapping neural representations of nearby attended locations and with the continuous nature of the attentional field. Recent fMRI work estimating the attentional field directly from visual cortical activity similarly supports the view that covert attention is represented as a spatially structured distribution rather than as a discrete categorical state (Bloem et al., 2025).

### Temporal stability and variability of the decoded attentional locus

The spatial structure of decoding errors is also relevant in light of evidence that covert spatial attention is intrinsically dynamic. Invasive recordings in non-human primates have shown substantial trial-by-trial fluctuations in the estimated attentional locus and demonstrated that the proximity of the decoded attentional spotlight to the behaviorally relevant location predicts perceptual performance (Astrand et al., 2016; De Sousa et al., 2021). Electrophysiological studies further suggest that attentional sampling exhibits rhythmic dynamics, particularly within alpha-and theta-frequency ranges (Fries, 2023; Gaillard et al., 2020; Gaillard and Ben Hamed, 2022). Recent human MEG decoding provides converging non-invasive evidence that the decoded attentional locus varies dynamically as a function of task demands (Mostafalu et al., 2026). Current MEG work indeed reports spatial decoding of covert attention together with task-dependent temporal fluctuations in the decoded representation.

The present fMRI results provide a complementary view at a much slower temporal scale. Cross-temporal decoding showed that classifiers trained around the late cue-to-target period generalized across several successive BOLD samples. This indicates that the multivoxel spatial code available to the classifier was sufficiently similar over time to support temporal generalization. At the same time, our within-trial analysis revealed that trials correctly decoded at target onset were not necessarily assigned to the same spatial location at earlier time points. The probability of recovering the cued location increased as target onset approached, a pattern consistent with a progressive stabilization or strengthening of the task-relevant spatial representation.

These observations should nevertheless be interpreted within the temporal limitations of fMRI. The BOLD response integrates neural activity over several seconds and successive fMRI samples are strongly shaped by the hemodynamic response and temporal autocorrelation. Consequently, broad cross-temporal generalization cannot establish that the underlying neuronal representation is stationary, nor can the present data resolve the rapid attentional fluctuations reported using electrophysiological recordings. Rather, our results indicate that these faster dynamics are embedded within a spatial representation that remains sufficiently consistent at the hemodynamic scale to support single-trial decoding. In this respect, fMRI and electrophysiological approaches provide complementary information: EEG and MEG can characterize rapid temporal fluctuations of attentional selection, whereas fMRI provides access to its spatially resolved cortical representation.

### Occipital cortex contains most of the fine-grained spatial information available to the decoder

Regional analyses revealed a marked difference between occipital and parietal contributions to decoding. Restricting feature selection to occipital cortex reproduced most of the performance obtained with whole-brain informative voxels, whereas decoding based exclusively on parietal voxels remained close to chance for the eight-location classification. Adding parietal voxels to occipital voxels provided only limited improvement. These results indicate that the information required for fine-grained localization of the attentional locus is predominantly expressed in occipital activity.

This finding is consistent with the retinotopic organization of the visual system. Spatial attention modulates activity throughout visual cortex in a location-specific manner, including in the absence of changes in retinal stimulation (Brefczynski and DeYoe, 1999; Datta and DeYoe, 2009; Kastner et al., 1999, 1998) Retinotopic mapping studies have demonstrated highly ordered representations of visual space across occipital visual areas (Arcaro et al., 2009; Warnking et al., 2002), providing an anatomical substrate from which multivariate classifiers can recover spatially precise attentional information. The spatial organization of the attention-selective voxels observed here was consistent with this known retinotopic architecture. Recent decoding studies reinforce this interpretation, showing that anticipatory attention can be decoded throughout visual cortex (Meyyappan et al., 2025) and that visuocortical activity can be used to reconstruct the spatial profile of the attentional field itself (Bloem et al., 2025).

The relatively weak eight-class decoding from parietal cortex should not be interpreted as evidence for a limited role of parietal cortex in spatial attention. A large body of work implicates dorsal frontoparietal regions in the endogenous control and allocation of spatial attention (Boshra and Kastner, 2022; Corbetta et al., 2008a; Corbetta and Shulman, 2002a). Contemporary models propose that control signals originating in the dorsal attention network bias spatially organized sensory representations in visual cortex. Consistent with this framework, our conventional univariate analyses showed robust recruitment of parietal as well as occipital and frontal regions during the attentional task. Moreover, attention-selective parietal voxels showed a contralateral organization despite the limited ability of parietal activity alone to distinguish the eight individual locations.

One interpretation is therefore that parietal cortex carries spatial information at a larger level, or in a representational format less directly accessible to the present eight-class linear decoder, whereas visual cortex expresses spatial attentional modulation within an explicitly topographic sensory map. This distinction is compatible with a functional architecture in which frontoparietal regions contribute to attentional control while visual areas provide a spatially precise expression of the resulting top-down bias. Importantly, however, the present data do not establish a causal direction of information flow between these regions, and the apparent difference in spatial scale should be tested directly in future studies.

### Visual-field asymmetries in the decoded representation of attention

The spatial organization of decoding revealed a more limited upper–lower visual-field asymmetry than initially observed. Rather than reflecting a general advantage for lower-field targets, the difference was restricted to a specific angular offset: prediction rates were significantly higher for upper-than lower-field targets at −π/4 in the whole-brain decoder, whereas the difference at the cued location itself did not reach significance. This spatially specific effect is nevertheless consistent with previous reports showing that visual processing and spatial attention can differ between the upper and lower visual fields (Carrasco et al., 2004; Montaser-Kouhsari and Carrasco, 2009; Zénon et al., 2009) although the present data do not support a simple overall lower-field advantage.

Our results therefore suggest that visual-field elevation can modulate the spatial structure of decoding errors, but that this anisotropy is subtle and depends on angular distance from the attended location. The underlying mechanism cannot be determined from the present data and could reflect differences in the precision, strength, cortical extent, or signal-to-noise ratio of the neural representation across visual-field locations. Moreover, no systematic difference was observed between horizontal and diagonal locations, suggesting that the spatial anisotropy detected here was specifically related to upper–lower field organization rather than to angular position more generally.

### Stability across training blocks and scanning sessions

A central requirement for participant-specific decoding applications is that classifiers remain reliable across repeated measurements. Decoding performance increased as more training blocks were included, with the largest improvements occurring during the initial accumulation of training data and comparatively limited gains after approximately 10–12 blocks. Importantly, decoding remained substantially above chance across four sessions acquired over two weeks, without evidence for a systematic deterioration in performance.

This reproducibility is encouraging for applications in which a participant-specific decoder must be calibrated before being used repeatedly. Rather than requiring a new classifier to be constructed for every session, a sufficiently informative training dataset may provide a stable basis for subsequent decoding. The substantial inter-individual variability observed here nevertheless indicates that calibration requirements are unlikely to be identical across participants. Future work should therefore determine prospectively how much training data are required to reach a predefined level of out-of-sample performance and whether calibration can be further improved through individualized ROI definition, acquisition parameters, or adaptive feature-selection procedures.

These considerations are particularly relevant to decoded neurofeedback and closed-loop fMRI. Real-time fMRI approaches increasingly use multivariate and machine-learning methods to target distributed neural representations rather than the activity of a single region (Cortese et al., 2017; Taschereau-Dumouchel et al., 2022; Weiskopf, 2012). Recent reviews emphasize both the potential of such approaches and the importance of rigorous participant-specific calibration, validation, and control conditions before clinical translation. The spatially resolved decoder developed here could ultimately provide a signal for closed-loop paradigms in which feedback depends on where attention is allocated rather than simply on a binary attended/unattended state. Such an approach could complement previous neurofeedback studies targeting attentional or higher-order cognitive states (Cortese et al., 2017; deBettencourt et al., 2015; Loriette et al., 2022, 2021).

### Limitations and future directions

Several limitations should be considered. First, although fMRI provides substantially greater spatial specificity than non-invasive electrophysiological methods, its temporal resolution is intrinsically constrained by the hemodynamic response. The present temporal analyses therefore characterize the evolution of the BOLD representation of attention rather than the instantaneous neuronal position of the attentional spotlight. In particular, the increase in decoding observed after target presentation is likely to contain both attentional information and target-evoked visual activity. The decoding observed at target onset, before the delayed hemodynamic response to target presentation, therefore provides a particularly important demonstration that spatial information was already present during anticipatory attention.

Second, decoding analyses were primarily based on behaviorally correct trials. The present findings therefore establish the information available when participants successfully performed the attentional task but do not determine how the neural representation differs on behavioral-error trials. The relationship between trial-wise decoding strength, decoded spatial error, and perceptual performance represents an important direction for future work, particularly given previous invasive and fMRI evidence linking attentional representations to behavior (Astrand et al., 2016; De Sousa et al., 2021; Meyyappan et al., 2025).

Third, although the present sample is sufficient to demonstrate robust group-level decoding and within-participant reproducibility, the marked variability between participants highlights the need for replication in larger samples. Understanding the anatomical, functional, and methodological factors underlying this variability will be particularly important for applications requiring reliable individual-level estimates.

Finally, spatially resolved decoding of covert attention requires careful separation of attentional signals from oculomotor activity. Covert attention and eye-movement systems are closely related, and even during fixation the direction of microsaccades can covary with spatial attention (Liu et al., 2022). Importantly, attention-related neural modulation can also occur in the absence of corresponding microsaccades, indicating that the two phenomena are related but not equivalent. In the present paradigm, gaze was monitored continuously and trials containing deviations outside the predefined fixation window were discarded and replayed. Quantitative characterization of residual fixation biases and microsaccadic activity would provide an additional test of the extent to which fine-grained decoding is independent of subtle oculomotor signals.

## Conclusion

In conclusion, the present results show that the locus of covert spatial attention can be recovered from participant-specific 3T fMRI activity with substantially greater spatial granularity than a simple hemifield or quadrant distinction. The information exploited by the decoder is spatially structured, predominantly expressed in occipital cortex, and reproducible across scanning sessions. At the same time, trial-wise decoding reveals variability in the estimated attentional locus that is compatible with the dynamic nature of spatial attention. These findings bridge fine-grained studies of attentional representations with participant-specific fMRI decoding and provide a foundation for future closed-loop approaches in which spatially resolved attentional states can be tracked and ultimately targeted through neurofeedback.

## Ethics declarations

The experimental protocol was approved by the French Ethic Committee (Comité de protection des personnes, #2018-72, ID RCB: 2017-A03612-51), and complied with the Declaration of Helsinki.

## Data and code availability

Data and code are available from the corresponding authors upon submission and approval of a formal project outline describing the intended use of the data.

## Author contribution

Conceptualization, C.L. and S.B.H.; Data Acquisition, C.L., C.D.S., S.D.; Methodology, C.L., S.D., S.C., F.L., D.I. and S.B.H.; Investigation, C.L., S.D. and S.B.H.; Writing – Original Draft, C.L., S.D.; Writing – Review & Editing, S.D., C.L., C.D.S., S.C., F.L., D.I. and S.B.H.; Funding Acquisition, S.B.H.; Supervision, S.B.H.

## Declaration of AI use

Generative AI tools were used to assist with language correction during manuscript drafting and with writing/debugging of analysis code. All outputs were reviewed and verified by the authors, who take full responsibility for the manuscript’s content.

## Supplementary material

**Supplementary Figure S1.**
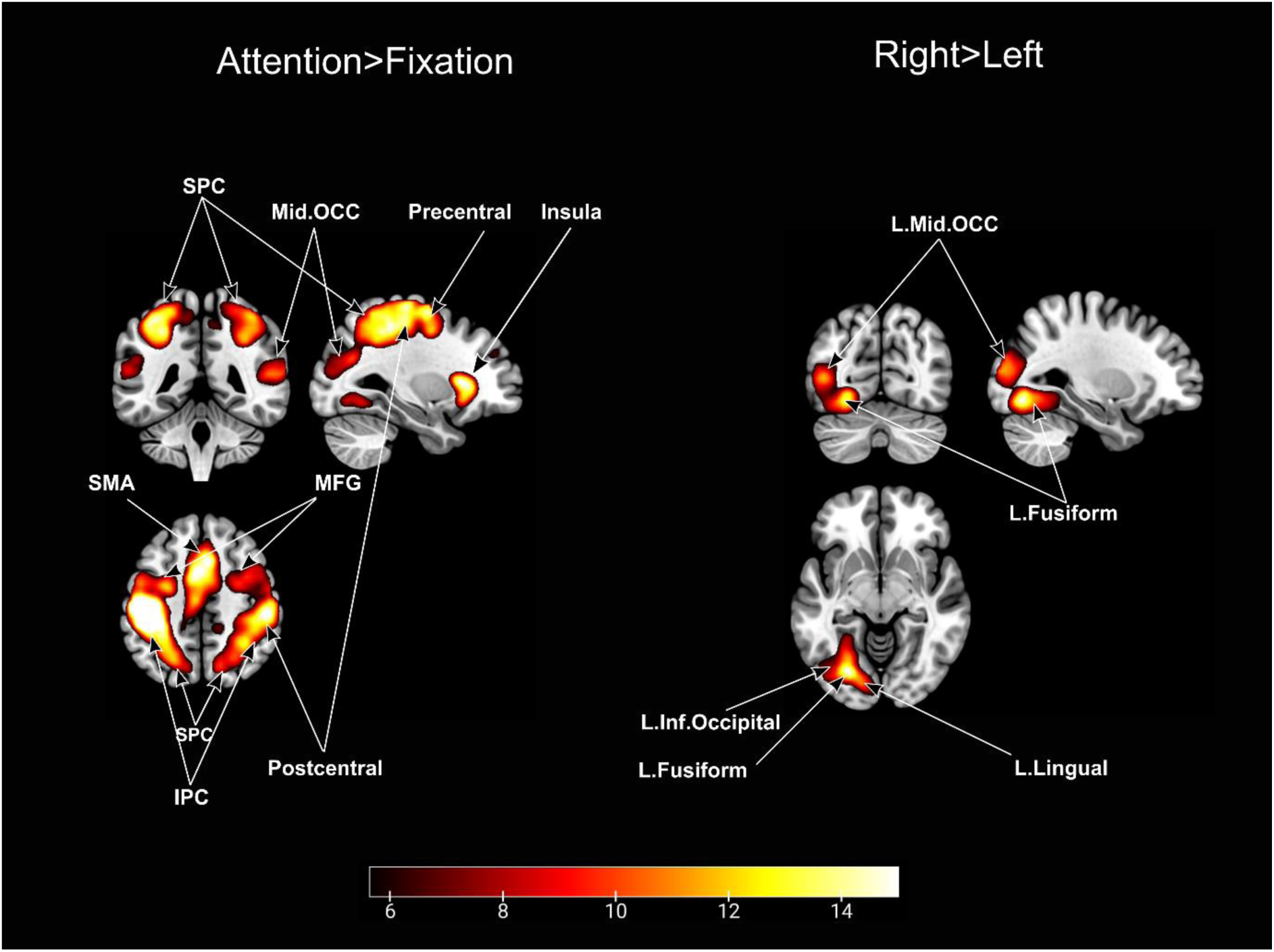
Whole brain attention-related brain activations. SPM activation maps associated with visuospatial attention. Group-level statistical maps are shown for the Attention > Fixation contrast (left) and the Right > Left attentional contrast (right). Attention relative to fixation recruited a bilateral network encompassing occipital, parietal, frontal, and sensorimotor regions. The Right > Left contrast predominantly involved left occipito-temporal regions, including the middle and inferior occipital cortices, fusiform gyrus, and lingual gyrus. Only clusters surviving family-wise error correction at p < 0.05 (FWE-corrected) are displayed. The color scale indicates statistical T values.

**Figure S2.**
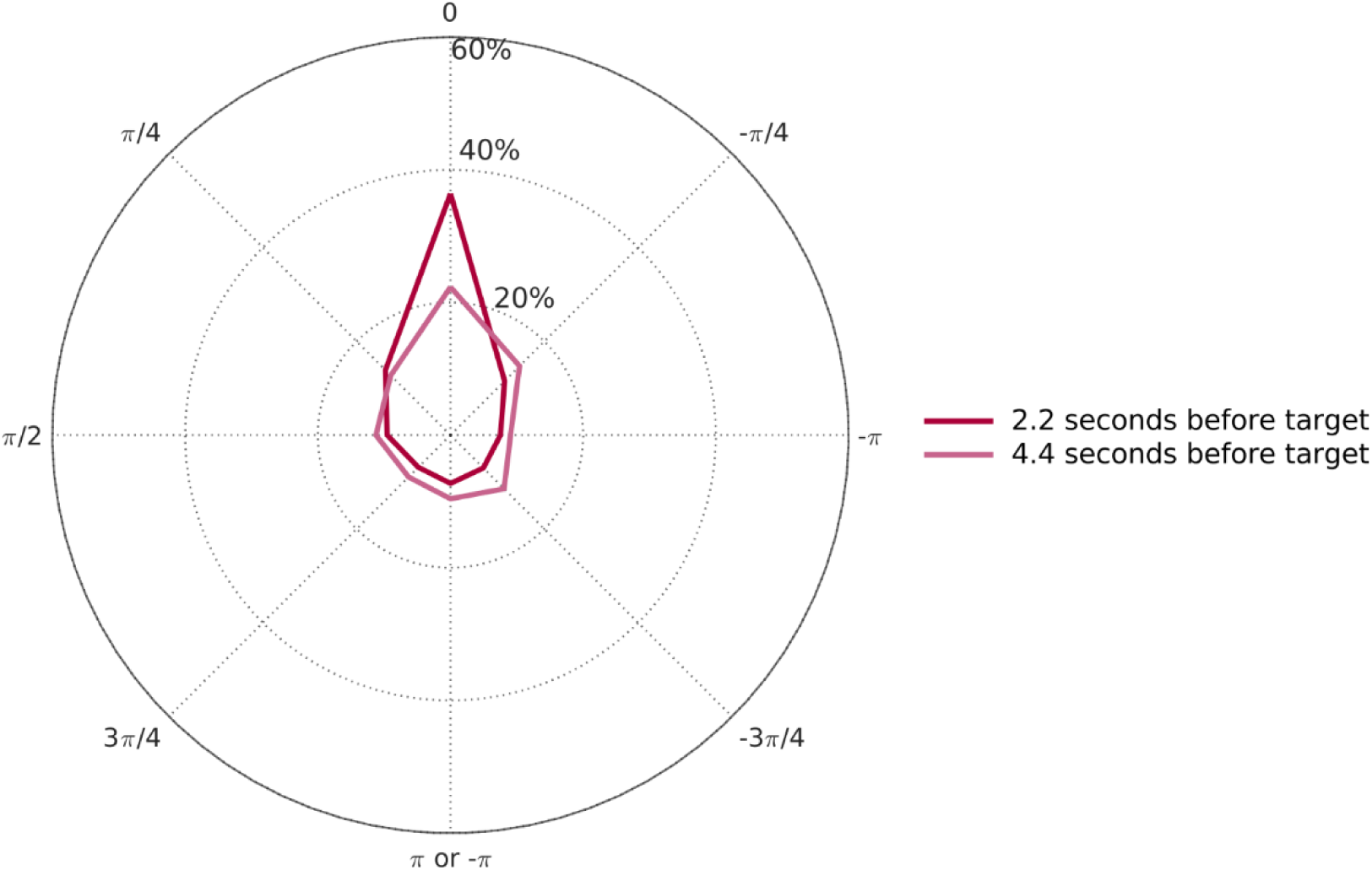
Temporal stability of spatial-attention decoding before target onset. The polar plot shows the percentage of decoded positions relative to the cued location at 2.2 s (dark red) and 4.4 s (pink) before target onset, considering only trials correctly classified at target onset. A value of 0 indicates the cued position, whereas increasing angular distances reflect decoding errors toward other locations. This analysis assesses the trial-by-trial stability of the decoded attentional position over time.

**Supplementary Figure S3.**
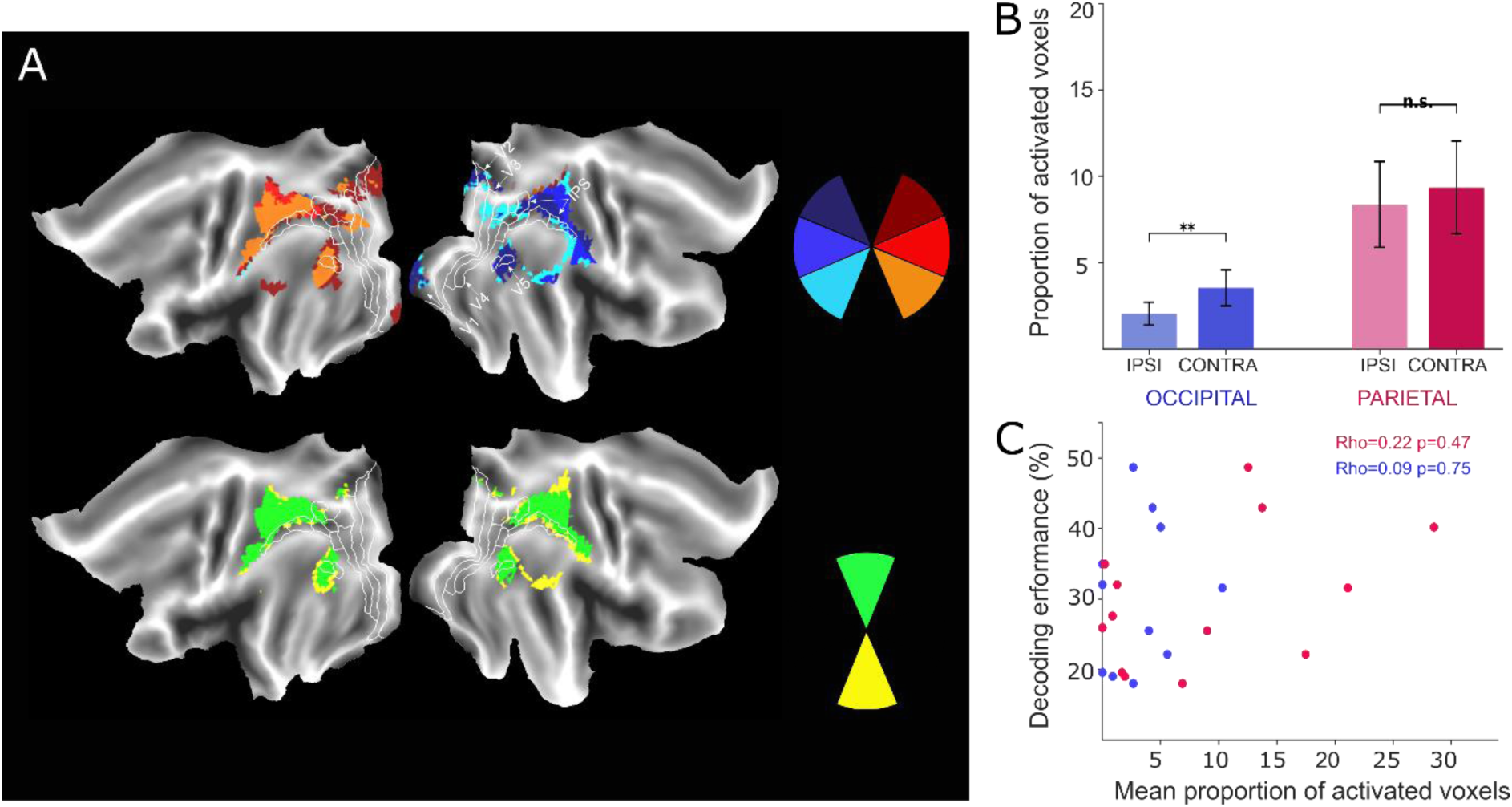
Spatial organization of task-activated voxels across attended locations. (A)Statistical parametric maps were computed independently for each of the eight cued positions by contrasting attention-related activity with the fixation baseline. Only significantly activated voxels were retained (p < 0.05, FWE-corrected). The flat maps on the left show the spatial distribution of activated voxels associated with the different attended locations; the six lateralized positions are shown in the upper maps, whereas the two positions located along the vertical meridian are shown separately below. (B) The upper-right panel shows the mean proportion of activated voxels located ipsilateral and contralateral to the attended hemifield within occipital and parietal cortices. (C) the relationship between decoding performance and the mean proportion of activated voxels across participants. No significant correlation was observed in either occipital.

